# Decoding Condition-Specific Cellular Crosstalk in Spatial Omics via Bilinear Edge Classification

**DOI:** 10.64898/2026.05.03.722470

**Authors:** Jonathan Karin, Roy Friedman, Mor Nitzan

**Affiliations:** School of Computer Science & Engineering, The Hebrew University of Jerusalem; Racah Institute of Physics, The Hebrew University of Jerusalem; Faculty of Medicine, The Hebrew University of Jerusalem

## Abstract

Tissues are multicellular structured communities whose function emerges from a combination of individual cellular characteristics along with their corresponding spatial configuration, affecting their interactions and response patterns. During processes such as disease progression or aging, tissues can undergo structural reorganization, including changes in co-localization of different cell types, assembly or destruction of functional niches, and disruption of intercellular communication axes. Such changes can manifest primarily in the spatial reorganization of cells rather than in the transcriptional states of individual cells. While computational tools for spatial transcriptomics have made significant progress in characterizing tissue architecture, most approaches for characterizing changes in tissue states across biological conditions operate at the level of individual cells or rely on discrete cell type labels, thus limiting the ability to detect coordinated transcriptional changes between neighboring cells that distinguish one condition from another. We present Casei, an interpretable bilinear edge classification framework comparing graphs across conditions, which directly models condition-specific cell-cell interactions in spatial omics data by focusing on interactions (edges), rather than cells (nodes), as the fundamental unit of inference. To capture such condition-specific signals, we leverage a model whose inductive bias aligns with cellular interactions through coordinated gene-gene relationships of neighboring cells, with learned weights that are directly interpretable as condition-specific gene-pair contributions. Casei enables the discovery of condition-associated multicellular interactions and spatial expression programs, and characterizes the loss of multicellular function and structure. We evaluate Casei on both controlled semi-synthetic spatial perturbations, where it outperforms node-level, edge-level, and graph neural network baselines, as well as on three real-world spatial omics datasets, where Casei enabled extracting underlying cell-cell interactions and interaction-informative genes and cells. Applied to mammalian liver fibrosis, atherosclerosis, and brain aging, Casei reveals biologically meaningful spatial reorganization, including the shift from endothelial-to macrophage-dominated networks in atherosclerotic plaques, disruption of hepatocyte zonation in fibrosis, and oligodendrocyte-microglia crosstalk in aging white matter.

## 1 Introduction

Characterizing the transition of tissues from healthy functioning units to collective malfunction, as can happen during disease progression, could reveal early biomarkers for tissue-level failure and routes for potential intervention strategies for delaying or reversing such processes [1, 2]. Some aspects of such tissue-level transitions manifest at the level of individual cell states; gene expression programs shift, cell type proportions change, and cellular subpopulations emerge [3– 6]. Recent computational methods have made substantial progress in detecting these cell-level state shifts from single-cell transcriptomic data, for example, by using graph signal processing to estimate the likelihood of observing each cell under different experimental conditions (MELD [7]) or identifying disease-affected cell subpopulations by refining case-control labels (HiDDEN [8]).

Recently we have shown through Annotatability [9] that condition-associated biological states can be identified by analyzing the training dynamics of a neural network learning to classify the gene expression profiles of cells exposed to different conditions, e.g. healthy vs. diseased. Annotatability is based on the observation that cells whose expression profiles are strongly associated with a specific condition are classified correctly early and consistently throughout the training process, whereas cells whose expression is not associated with any of the labeled conditions receive unstable, fluctuating predictions across training epochs. Therefore, features extracted from the training dynamics of neural networks over a single-cell dataset can be used as a measure of condition-association at single-cell resolution [9].

However, some tissue-level physiological changes are not manifested merely in individual cellular states, but rather in the structural reorganization of the tissue, involving changes of adjacency structures of different cell types, assembly of functional niches, and disruption of intercellular communication pathways. From a machine learning perspective, this casts the problem as one comparing graphs across conditions and identifying differentiating edges. For example, in triple-negative breast cancer, the spatial distribution of CD8 T cells within the tumor microenvironment differs between treatment responders and non-responders [10], and in colorectal cancer, tumors reorganize into spatially defined multicellular neighborhoods with characteristic cell compositions and local interactions that correlate with malignancy and clinical outcome [11].

Characterization of such cellular reorganization can be probed by spatial transcriptomics technologies [12, 13], providing spatially-aware gene expression measurements. While spatial reorganization can be accessible via co-localization statistics between different cell types [14], this process requires discrete cell type labels and is therefore sensitive to the granularity of annotation. Since cells generally exhibit continuous variability in their transcriptional states, categorical co-localization counts cannot capture coordinated changes in gene expression between neighboring cells that are more subtle than cell type-level co-localization patterns.

As a more elaborate alternative, detecting condition-associated spatial phenotypes can be done using a machine learning classifier, yet this poses an inductive bias problem; While such classifiers learn to distinguish between conditions based on patterns in the input data, the patterns that can be revealed by a classifier depend on assumptions encoded in its architecture regarding the signal of interest. Therefore, identifying condition-specific multicellular spatial organization relies on encoded assumptions about the target forms of spatial phenotypes. For example, patterns learned by models that represent each cell independently reflect condition-associated changes that manifest in individual cell states, while patterns learned by models that aggregate features of neighboring cells, like graph neural networks, reflect condition-associated signals that lie in combined properties of spatial cellular neighborhoods. Therefore, the architecture of the model does not merely influence classification accuracy, but determines which types of biological variations can be extracted from the data. Interpretability is equally important, where biological discovery can be facilitated when the genes driving an observed difference emerge directly from a classifier’s learned weights, rather than inferred after the fact through post-hoc analysis.

Here we focus on condition-specific tissue reorganization that is reflected in coordinated gene-gene co-expression patterns between neighboring cells. Therefore, we construct a model whose inductive bias aligns with the co-expression relationships between neighboring cells, by shifting the object of inference from cells to the interactions between them. Therefore, rather than asking which cells are condition-specific, we ask: which pairwise relationships between neighboring cells, namely which *edges* in the cellular spatial proximity graph (where nodes represent cells and edges connect spatially-proximal cells [15]), are condition-specific? We present Ca-sei (Condition-Associated Spatial Edge Inference), a bilinear and interpretable-by-construction classifier that explicitly models coordinated gene-gene interactions between neighboring cells (Fig. 1) motivated by an energy-based model for pairwise interactions (Methods 4.5). For each condition, Casei learns a separate low-rank interaction matrix encoding the coordinated gene expression patterns between neighboring cells that are predictive of that condition. Edges are then ranked based on features extracted from the model’s training dynamics and filtered to retain only cell-cell interactions most robustly associated with each condition. This results in a condition-adjusted cellular proximity graph, that can be plugged directly into any existing graph-based spatial analysis tool, enabling condition-associated spatial analysis.

**Figure 1:**
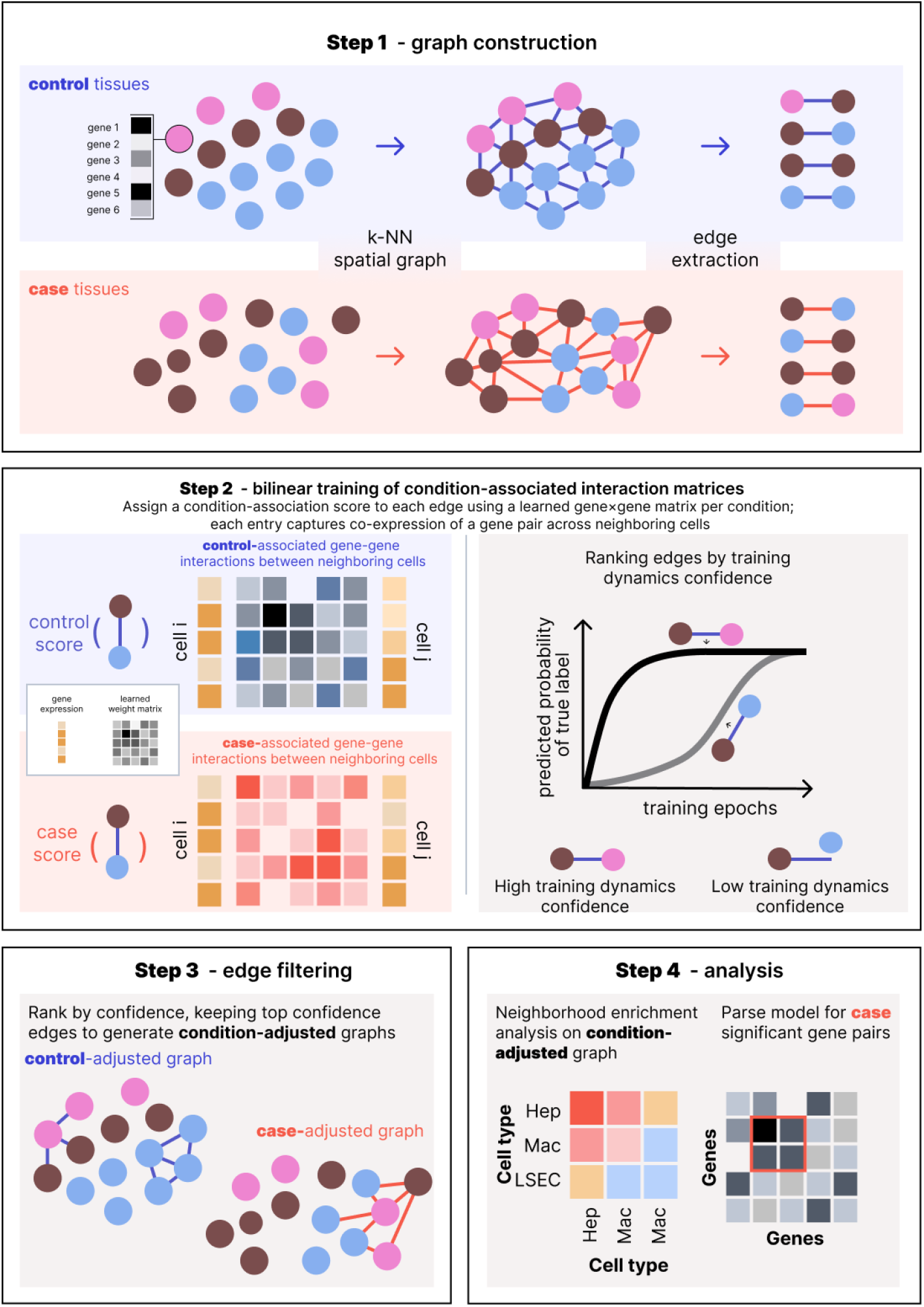
Overview of Casei, a framework for the identification of condition-specific cell-cell interactions in spatially-informed gene expression data, using a training dynamics-based bilinear classifier, illustrating the pipeline from spatial graph construction and bilinear model training to edge filtering and downstream analysis.

We evaluate Casei on both semi-synthetic spatial perturbations, where it outperforms seven node-level, edge-level, and graph neural network baselines, and on three spatial transcriptomics datasets, where it reveals biologically meaningful spatial reorganization patterns. Including the shift from endothelial-dominated to macrophage-dominated networks in atherosclerotic plaques, the disruption of hepatocyte zonation and loss of Kupffer cell interactions in liver fibrosis, and oligodendrocyte-microglia crosstalk in aging brain white matter. Beyond identifying signal-specific cell-cell interactions, Casei can be used to extract condition-associated enriched genes and infer condition-specific spatial expression programs.

## 2 Results

### 2.1 Overview of Casei: a confidence-based cell-cell edge classification framework

Casei takes as input a collection of *N* spatial transcriptomics samples 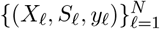, where 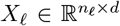 is the gene expression matrix of sample *l* with *n*_*l*_ cells and *d* genes 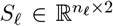 are the associated physical coordinates for each cell (or pseudo-cell) which we consider to be 2-dimensional (yet can be generalized to 1 or 3 dimensions given other measurement types), and *y*_*l*_ *∈* {1, …, *C*} is a condition label for each sample (*e*.*g*. healthy or diseased). First, we construct *N k*-nearest-neighbor (*k*-NN) spatial proximity graphs 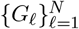 based on the spatial coordinates *S*_*l*_ for each sample independently, where each node *i* in *G*_*l*_ corresponds to a cell with gene expression profile **x**_*i*_ ∈ ℝ^*d*^ from the corresponding gene expression matrix *X*_*l*_, and each edge (*i, j*) connects two spatially proximal cells (**x**_*i*_, **x**_*j*_), inheriting the condition label of their sample (Fig. 1, Step 1).

#### Gene-Gene Interaction Modeling via a Bilinear Architecture

For each condition *c*, we learn a pairwise cell interaction representation *M*_*c*_, which is modeled as symmetric positive semi-definite and bilinear through the matrix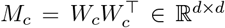, where *W*_*c*_ ∈ ℝ^*d×r*^ and *r ≪ d*, in turn making *M*_*c*_ low-rank, to keep computation tractable. This parameterization explicitly captures coordinated gene-gene co-expression between neighboring cells while remaining interpretable; (*M*_*c*_)_*k,m*_ directly encodes the strength of coordinated expression between gene *k* in one cell and gene *m* in its neighboring cell, thus enabling direct identification of gene pairs that drive condition-specific interactions. For each edge (*i, j*) in the spatial proximity graph, we define the *interaction score* for condition *c* between neighboring cells with gene-expression profiles **x**_*i*_, **x**_*j*_ *∈* ℝ^*d*^ as:

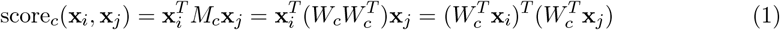

The interaction matrices *M*_*c*_ are then optimized as classifiers between the different conditions using the standard cross-entropy classification loss:

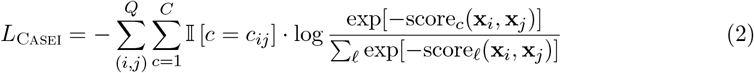

where *Q* is the total number of edges over all *k*-NN spatial proximity graphs, *c*_*ij*_ *∈* { 1,*· · ·, C*} is the input condition label for each edge, and 𝕀 [*c* = *c*_*ij*_] equals 1 when *c* = *c*_*ij*_ and 0 otherwise (Fig. 1, Step 2). Training Casei in this way is theoretically grounded through energy-based modeling of pairwise interactions, where the accumulated interaction score for a whole tissue depicts the energy of the transcriptomic sample (Methods 4.5).

#### Confidence Quantification Through Training Dynamics Analysis

Not all cell-cell interactions, or edges, in a tissue are condition-associated. To identify proximal cell pairs whose co-expression patterns are associated with a specific condition, we exploit information that can be gained from the training dynamics of the classifier [9, 16]. Specifically, we train the bilinear classifier by minimizing the cross-entropy loss *L*_Casei_ via stochastic gradient descent, updating the weight matrices {*W*_*c*_} across *E* epochs. At each epoch *e*, the model assigns a probability *p*^(*e*)^(*c*_*ij*_ |**x**_*i*_, **x**_*j*_) to the input condition label of each edge (*i, j*). As training progresses, this probability tends to increase over epochs for edges that are predictive of the condition and therefore condition-associated, while remaining low or fluctuating for non-predictive edges [9, 16]. As proposed in Annotatability [9] in the context of individual cells, averaging the predictions across epochs, rather than using only the final model’s predictions, captures continuous biological variation masked by discrete labels. We therefore define the confidence of edge (*i, j*) as the mean predicted probability across all training epochs, *E*:

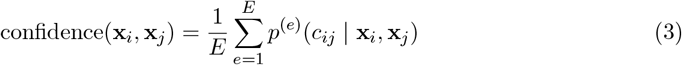

where *p*^(*e*)^(*c*_*ij*_ |**x**_*i*_, **x**_*j*_) is the prediction probability for the input condition label *c*_*ij*_ that the model assigns to the edge (*i, j*) at epoch *e*.

Finally, we construct a *condition-adjusted graph* (Fig. 1, Step 3; Methods 4.8), which is a subgraph of the per-sample spatial proximity graph, by retaining only high-confidence edges (Methods 4.2) to enable spatial signal-aware downstream analysis (Fig. 1, Step 4).

### 2.2 Bilinear edge classification accurately recovers diverse spatial reorganization patterns

We first evaluated the performance of Casei using mouse organogenesis seqFISH (sequential fluorescence in situ hybridization) data [17] which we perturbed in-silico. The dataset comprises spatial transcriptomics measurements of 19,416 cells, profiling 351 genes and spanning 22 annotated cell types across the developing brain, heart tube, gut tube, and neural tube [17]. The synthetic perturbations, or different cases, we used were designed to isolate different aspects of spatial tissue reorganization, and were each applied to a subregion of the tissue (Fig. 2a-e, Methods 4.6): **Case 1:** randomly shuffles cell positions within a predefined region; **Case 2:** replaces the expression profiles of a fraction *p*_enrich_ of cells within a predefined region with profiles sampled with replacement from a target cell type; **Case 3:** swaps the expression profiles of cells belonging to two predefined cell types within a predefined region; **Case 4:** enriches spatial co-localization of two predefined cell types by relocating cells of one type into the neighborhood of the other within a predefined region. These four types of perturbations alter the spatial organization of cells, while keeping the distribution of cell states in the tissue unchanged, which in turn allows to probe the sensitivity of different models to detecting different modes of spatial reorganization (see Methods 4.6 for full simulation details).

**Figure 2:**
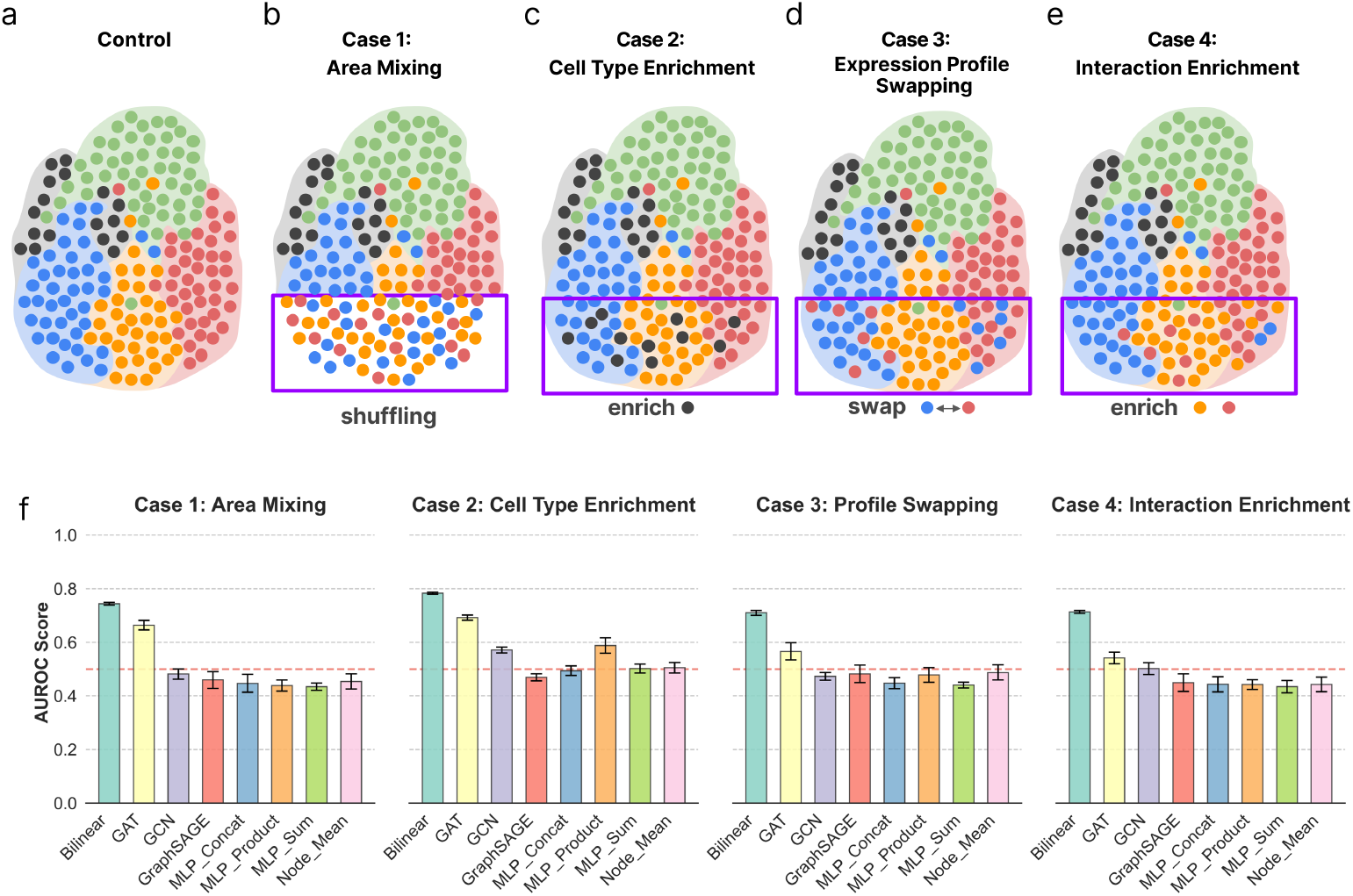
Evaluating identification of synthetic perturbations over spatial transcriptomics data. **(a-d)** Visualization of the synthetic perturbations setup based on a mouse organogenesis seqFISH dataset [17]. **(a)** Unperturbed control tissue, colored by cell type. **(b)** Case 1: Area mixing perturbation, where cells are shuffled locally. **(c)** Case 2: Cell type enrichment perturbation, where a specific cell type population is enriched locally. **(d)** Case 3: Cell type swapping perturbation, where the locations of cells of two cell types are exchanged within a local tissue region. **(e)** Case 4: Interaction enrichment perturbation, generating co-localization enrichment of pairs of cells of two specific cell types. **(f)** Benchmarking performance across the four perturbation scenarios. Barplots display the Area Under the ROC Curve (AUROC) for edge-level perturbation detection (n = 10 replicates). The proposed **Casei** (bilinear model) architecture is compared against a node-centric MLP baseline (**Node Mean**), three node-level graph neural network baselines (**GCN** [18], **GAT** [19], **GraphSAGE** [20]), and three edge-level MLP variants (**MLP_Sum, MLP_Concat, MLP_Product**). For node-level baselines (Node Mean, GCN, GAT, GraphSAGE), edge confidence is computed as the average confidence of the two incident node confidences. The dashed red line indicates performance of a random classifier (AUROC=0.5).

We frame the evaluation as an *edge classification* task between paired control (original data) and case (synthetically perturbed) tissues. Models were evaluated by their ability to identify (assign higher confidence) to perturbed edges, defined as edges with at least one incident cell that was synthetically perturbed. Edge classification was evaluated for each of the perturbation types using the AUROC score of edges ranked by assigned confidence relative to the ground truth set of perturbed edges (Methods 4.6), averaged over four independently generated synthetic replicate tissues.

We first note that any method operating at the cell level, such as Annotatability [9], MELD [7], or HiDDEN [8], would not be able to detect these spatial perturbations, since the global distribution of gene expression profiles does not change, by design, in the perturbed settings.

We therefore compared the predictions of spatial perturbations of Casei to seven architectural variants (see Methods 4.6), spanning two families: The first family consists of node-level baselines that score individual cells and derive edge confidence as the mean of the two incident nodes’ scores; *Node-Mean* uses a standard MLP applied independently to each cell. *GCN* [18], *GAT* [19], and *GraphSAGE* [20] are graph neural networks that propagate information along the spatial cellular proximity graph before classification; these architectures form the backbone of many recent spatial omics tools for tissue domain identification, batch integration, and cell-cell communication inference [21–25], and we include them as representative node-level spatial baselines. The second family consists of edge-level MLP baselines that aggregate the expression profiles of neighboring cell pairs before classification, via element-wise summation (*MLP-Sum*), concatenation (*MLP-Concat*), or element-wise multiplication (*MLP-Product*).

Casei outperforms all other baselines in identifying perturbed edges across all four spatial perturbation cases described above (Fig. 2e), as well as two additional cases, including tissue-wide spatial permutation of cells and localized cell type-specific ligand-receptor expression induction (see Methods 4.6, Supplementary Fig. 6).

### 2.3 Casei Reveals Reorganization of Microvascular Interaction Networks in Human Atherosclerosis

In healthy arteries endothelial cells line the vessels and maintain stable contacts with their neighbors [26]. However, this organization can break down in disease and transition to dense inflammatory neighborhoods dominated by macrophages [27]. Specifically, atherosclerosis is characterized by remodeling of the cellular architecture of the vessel wall, initiated by lipoproteins accumulating in the inner lining of the artery, which triggers an inflammatory response. This, in turn, leads to infiltration of macrophages and inward migration of smooth muscle cells, which together form the core of growing plaques [28].

We studied the spatial cellular reorganization in atherosclerosis using Casei through the analysis of an atherosclerosis single-cell resolution Xenium spatial transcriptomics dataset [29], comprised of 12 atherosclerotic plaque samples and 4 healthy control coronary artery samples (see Supplementary Note 8).

We trained Casei to classify neighboring cell pairs as originating from healthy or plaque tissue (Fig. 3b, c). Then, we filtered the k-NN-spatial proximity graph constructed over all cells to retain a subgraph composed of the top 5% of edges confidently classified as healthy, generating a *healthy-adjusted graph* that reflects cell-cell interactions associated with the healthy condition (Methods 4.8). In the same manner, we constructed a *plaque-adjusted graph* composed of the edges most confidently classified as associated with plaques. Applying neighborhood enrichment analysis to these condition-adjusted graphs, which can be implemented through Squidpy [15], then reveals which cell types tend to co-localize among condition-associated interactions. Specifically, applying neighborhood enrichment to the healthy-adjusted graph (Fig. 3b, d, g) reveals that the cellular interaction landscape in healthy tissue is dominated by endothelial-endothelial interactions (Fig. 3b, d, g), as well as interactions between endothelial cells (EC) with both vascular muscle cells (VSMC) and macrophages (EC-VSMCs and EC-macrophages, Fig. 3h). The enriched endothelial-endothelial interactions reflect the structure of healthy vessel walls, where endothelial cells form an intact lining (Fig. 3b, d), which is lost with the introduction of plaque. Furthermore, Casei can be used to extract gene pairs that are associated with condition-specific interactions, based on differences in the healthy-associated and disease-associated interaction matrices, which we call the *differential interaction matrix* (Methods 4.7). This analysis revealed ITLN1 (Intelectin-1) as a central hub in healthy-associated interactions, appearing in all top 10 gene-gene interaction pairs whose coordinated expression across neighboring cells is most strongly associated with the healthy condition (Fig. 3f). ITLN1 is an adipokine with vascular expression that exerts anti-inflammatory, anti-atherogenic effects, including reduction of macrophage foam cell formation and vascular inflammation [30]. ITLN1 coordinated expression with extracellular matrix components (*LUM, MMP12, MMP13*) and immune regulators (*CCR2, CD68, CCL18*), identified by Casei, point to a possible coordinating role at the interface of endothelial barrier integrity, matrix homeostasis, and baseline immune surveillance. In contrast, cellular neighborhood enrichment of the plaque-adjusted graph showed a shift toward macrophage-dominated neighborhoods (Supplementary Note 8, Fig. 3f, i). This spatial reorganization reflects the transition from vascular homeostasis to chronic inflammatory microenvironments that is characteristic of atherosclerosis [31, 32].

**Figure 3:**
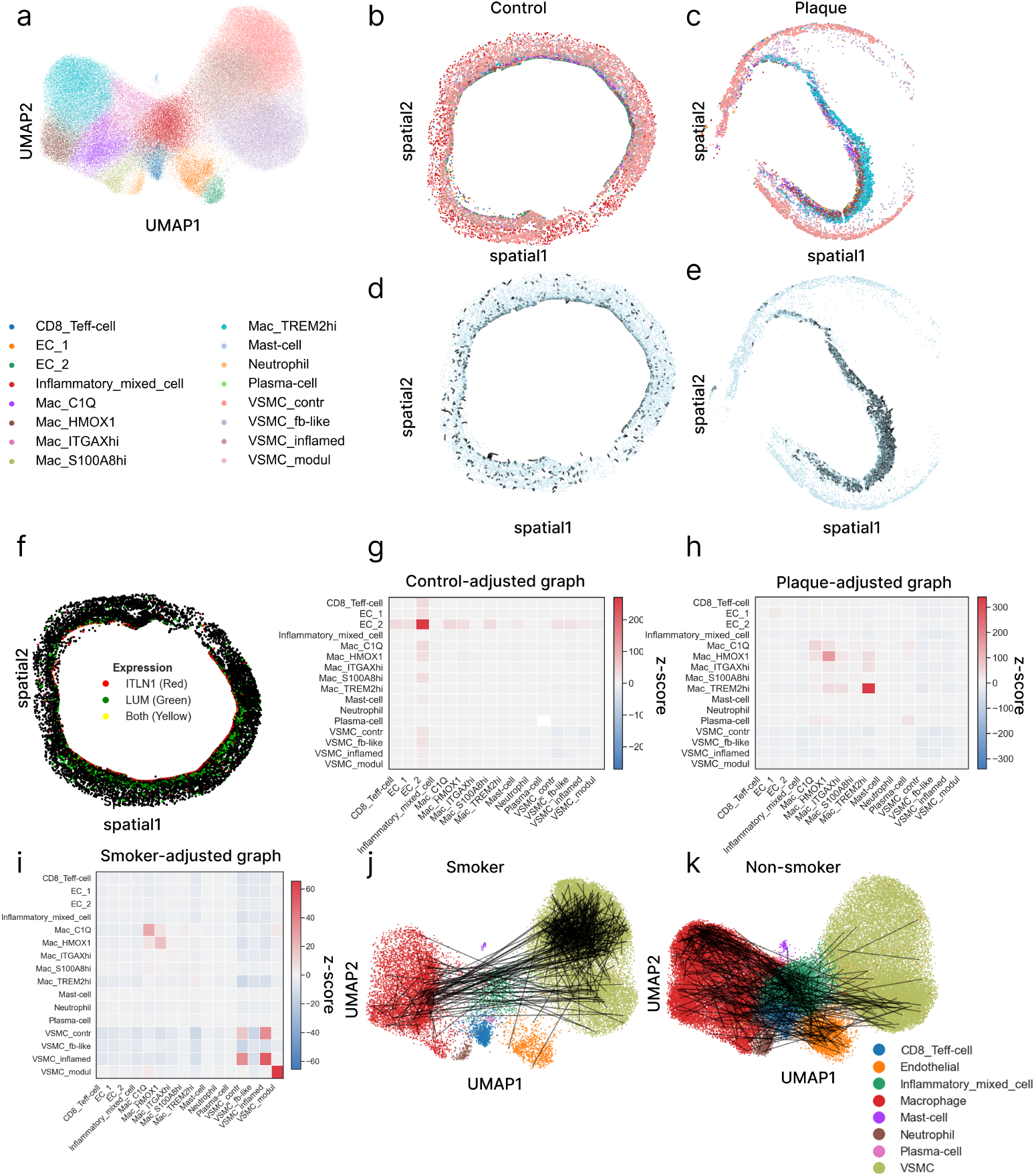
Casei reveals condition-associated spatial reorganization of cellular interaction networks in human atherosclerosis. **(a)** UMAP visualization of the atherosclerosis Xenium dataset [29] colored by cell type. **(b-e)** Spatial scatter plots of representative control (P4; left) and plaque (P7; right) patient samples showing healthy vessel vs. diseased tissue structure, colored by (b,c) cell type as in (a), and (d,e) training dynamics confidence scores, where cells are colored in light blue, and high-confidence condition-specific edges are colored in black. **(f)** Spatial scatter plot of a representative healthy tissue sample, where cells are colored by the expression level of ITLN1 (red) and LUM (green), the top-ranked gene pair based on the healthy-associated differential interaction matrix (see Methods 4.7). **(g-i)** Case-adjusted neighborhood enrichment heatmaps showing the shift from endothelial-dominated interactions in control tissue **(g)** to macrophage-dominated interactions in plaques **(h)**, and VSMC-centered interactions in smoker plaques **(i). (j-k)** Projection of the high-confidence interactions (black lines) onto the UMAP embedding (a) for smoker **(j)** and non-smoker **(k)** plaques, visualizing the connectivity shifts between macrophage-centered interactions in non-smoker plaques to VSMC-centered interactions in smoker plaques.

Next, based on patient-level smoking history for the plaque samples [29], we studied whether the spatial reorganization of plaques differs between smokers and non-smokers. We trained Casei to classify between smoker and non-smoker plaques, constructing both smoker-adjusted and non-smoker-adjusted spatial cellular graphs. Analysis of the smoker-adjusted graph revealed enrichment of VSMC-VSMC (specifically contractile and inflamed VSMCs) interactions in smoker plaques (Fig. 3i,j; Supplementary Note 8), while the non-smoker-adjusted graph showed enrichment of macrophage-macrophage interactions (Fig. 3k, Supplementary Fig. 10). These results suggest that smoking may be associated with a shift toward VSMC-centered spatial organization within plaques, relative to the macrophage-dominated interaction patterns observed in non-smoker plaques.

### 2.4 Inference of context-associated cell-cell interactions identifies the breakdown of functional tissue niches in liver fibrosis

Liver fibrosis is a progressive disease in which excessive extracellular matrix deposition leads to tissue remodeling and a breakdown of normal liver architecture [2]. Beyond changes in individual cell states, fibrosis disrupts the spatial organization of the tissue, dismantling functional niches. Identifying the healthy cell-cell interactions that are selectively lost during fibrosis could highlight functional niches whose disruption underlies disease progression. We applied Casei to spatial transcriptomics data that includes 3 healthy control liver tissue samples and 4 samples of fibrotic liver tissue (Fig. 4a-e), generated using MERFISH technology with 300 genes profiled per sample [2]. We trained Casei to discriminate between healthy and fibrotic liver tissues (Methods 4.2). We focused our downstream analysis on cell–cell interactions that characterize healthy liver tissue through the healthy-adjusted spatial graph (Methods 4.8), highlighting cellular interactions that are lost during fibrosis. Neighborhood enrichment analysis over the healthy-adjusted graph (Methods 4.8) revealed two dominant cell-cell interactions: hepatocyte-hepatocyte and hepatocyte-macrophage interactions (Fig. 4f, left panel).

**Figure 4:**
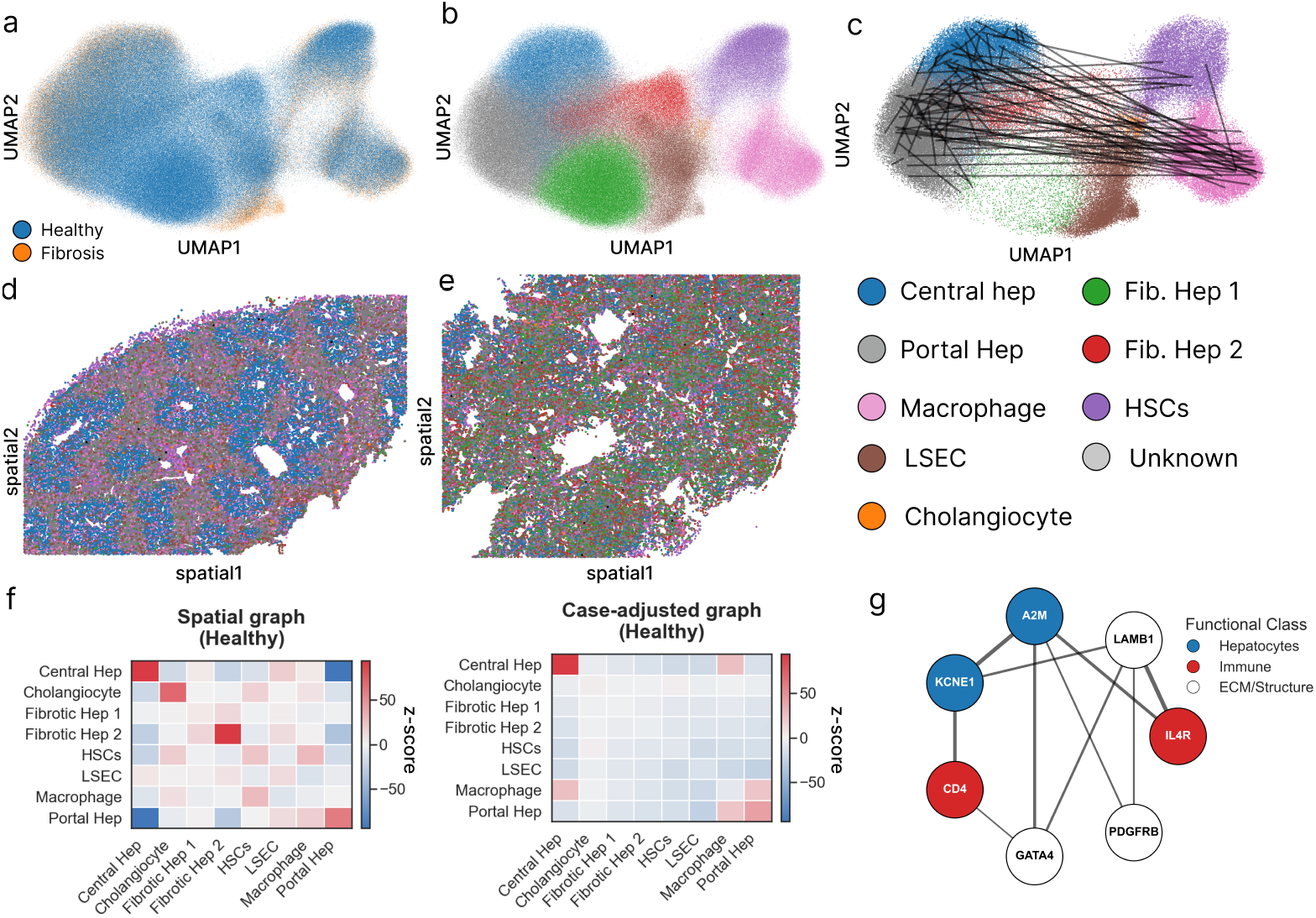
Application of Casei to liver fibrosis spatial transcriptomics reveals disease-associated interaction patterns. **(a**,**b)** UMAP visualization of the MERFISH liver dataset [2], colored by (a) disease status and (b) annotated cell types. **(c)** Projection of the healthy-associated cell-cell interactions learned by Casei (black lines) onto the UMAP embedding, colored as in (a) according to cell types. The cell-cell interaction patterns visualize the connectivity between the macrophage cluster and hepatocyte clusters. **(d-e)** Spatial scatter plots displaying cell type distribution in representative liver tissue sections from the diseased (fibrotic, b) and healthy (c) samples, colored according to cell types as in (a). **(f)** Neighborhood enrichment heatmap computed over the learned healthy-adjusted graph (left), versus the original spatial proximity graph over the cells in the healthy tissues. The healthy-adjusted graph contains two main interactions: hepatocyte-hepatocyte (reflecting preserved liver zonation) and hepatocytemacrophage (reflecting homeostatic crosstalk). **(g)** Network analysis of the top molecular drivers characterizing the healthy hepatocyte-macrophage niche. The diagram highlights the A2M-CD4 and A2M-IL4R axes as key mediators of the interaction between hepatocytes (blue) and immune cells (red), anchored by structural ECM components (grey).

The hepatocyte-hepatocyte interactions recovered by Casei in the healthy-adjusted spatial graph (Fig. 4c,f) reflect organized functional hepatocyte layers, where cells are spatially *zonated* and perform distinct metabolic tasks based on their spatial location along the central-portal lobule axis [33–35]. These interactions are lost in fibrosis by scar tissue and vascular changes, disrupting the spatial gradients of hepatocyte zonation [36].

Similarly, hepatocyte-macrophage interactions enriched in the healthy-adjusted graph (Fig. 4c, f) reflect homeostatic crosstalk between hepatocytes and Kupffer cells, resident liver macrophages, which reside within the sinusoids in close contact with hepatocytes and regulate metabolism and immune defense [37]. The absence of this specific interaction signature in the fibrotic case may reflect the disruption of the sinusoidal niche, as during fibrosis, resident Kupffer cells are often depleted or replaced by infiltrating inflammatory macrophages, severing the normal spatial communication required for liver stability [38]. The hepatocyte-macrophage cellular interaction was not identified by cell-level quantification using either MELD [7] or Annotatability [9] (Sup. Fig. 7, Methods 8).

Next, we constructed a network of genes, where gene pairs are differentially connected according to their condition-associated co-expression patterns (Methods 4.7; Fig. 4g, Supplementary Table 1). Those can be divided into three functional classes: hepatocytes-associated genes (e.g. *A2M, KCNE1* [39, 40]), *immune markers (e*.*g. CD4, IL4R*), and extracellular matrix (ECM)/structural components (e.g. *LAMB1, GATA4, PDGFRB* [41]) whose pairwise interactions recapitulate the healthy hepatic niche (Fig. 4f). Specifically, the gene-gene interaction network exhibits strong interaction strengths (namely large amplitudes within the differential interaction matrix, reflecting strongly coordinated expression across neighboring cells; Methods 4.7) for pairs of hepatocyte-hepatocyte coordinated gene expression within the organized hepatocyte layers; hepatocyte-ECM and immune-ECM pairs of genes reflecting the structural organization of the perisinusoidal ‘Space of Disse’ area in the liver, required for hepatocyte-endothelium contact [41, 42]; and hepatocyte-immune pairs of genes capturing the homeostatic crosstalk between hepatocytes and resident Kupffer cells described above. Together, these gene pairs paint a coherent molecular picture of the healthy liver niche that is dismantled in fibrosis - that is, the crosstalk between hepatocytes with other hepatocytes, ECM and immune cells is disrupted during liver fibrosis.

### 2.5 Casei Identifies Region-Specific Rewiring of Glial Interaction Networks in the Aging Mouse Brain

We next studied spatial reorganization of cellular interactions along aging through the analysis of a spatial transcriptomics brain atlas of young (3.4-5.4 months) and aged (28.5-34.5 months) mice [1] (Fig. 5a, c). After training Casei to distinguish aged from young brain tissue, we constructed young-adjusted and aged-adjusted cellular proximity graphs (Methods 4.8). Analysis of these graphs revealed that aging-associated cellular interactions were not spread uniformly across the brain, but were instead concentrated in specific anatomical regions. To infer which regions are most affected by aging-associated interaction rewiring, we identified those with a high proportion of cells participating in high-confidence edges (Fig. 5d), and characterized them by their cell type composition (Fig. 5b) and neighborhood enrichment patterns (Fig. 5f).

**Figure 5:**
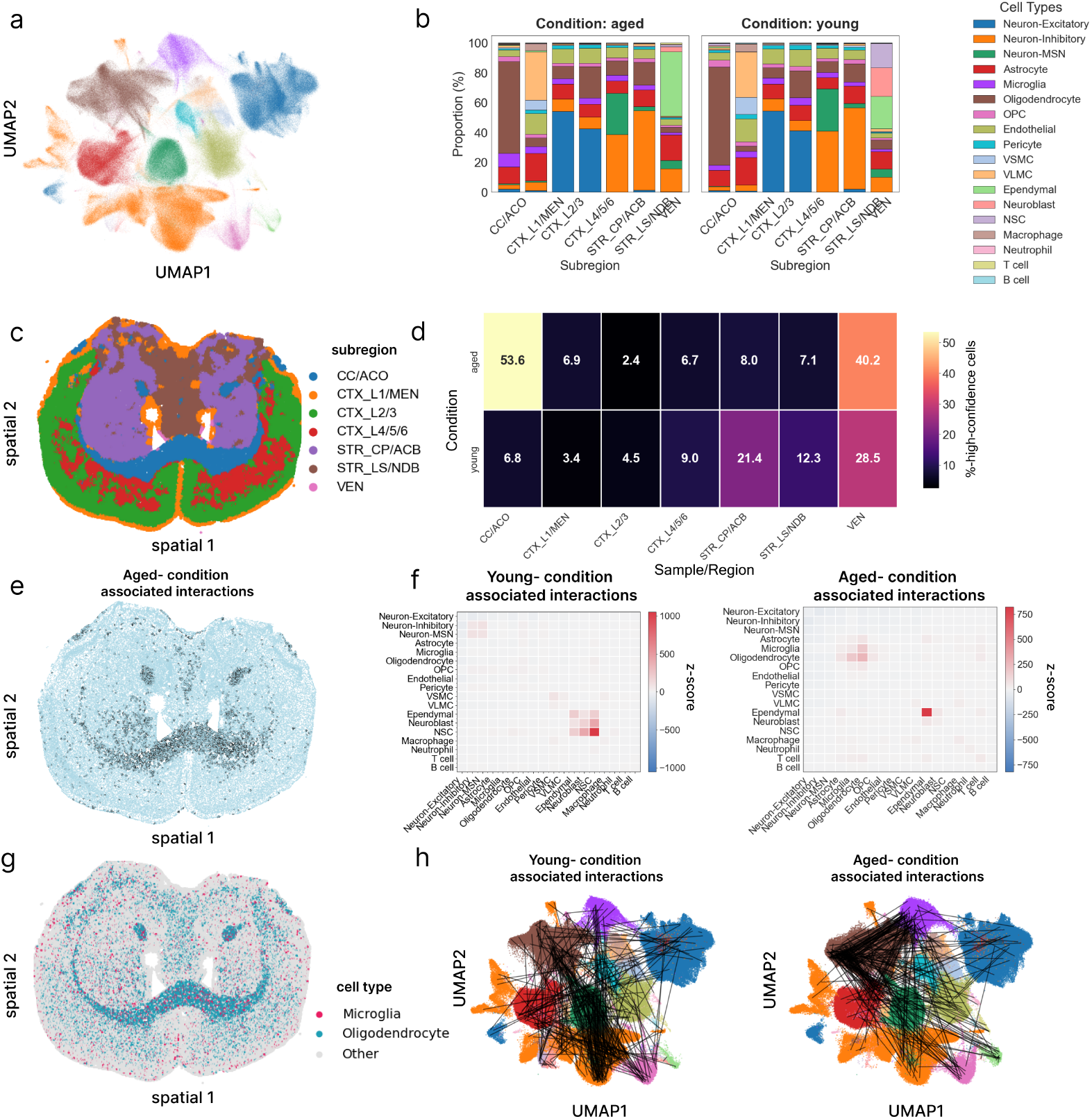
Condition-associated interactions in the aging mouse brain. **(a)** UMAP visualization of the mice brain MERFISH dataset colored by cell type [1]. **(b)** Cell type proportions in young vs. aged mice according to subregions. **(c)** Spatial scatter plot visualization of a coronal brain section of aged mouse (age 32.6) colored by subregion mapping. **(d)** Proportions of cells with at least one incident edge included in the condition-adjusted cellular proximity graph computed by Casei for aged (top) versus young (bottom) mice, per subregion. **(e)** Spatial scatter plot, as in (c), where cells are colored in light blue and high confidence edges are colored in black. **(f)** Neighborhood enrichment heatmap computed over the learned young-adjusted graph (left) and aged-adjusted graph (right). **(g)** Spatial scatter plot, as in (c), colored by cell types (microglia: red, oligodendrocytes: blue, others: gray). **(h)** Projection of a representative sample of 500 high-confidence interactions (black lines) onto the UMAP embedding for young-associated (left) and aged-associated (right) edges, visualizing the connectivity shifts from NSC-neuroblast-ependymal interactions in young tissue to oligodendrocyte-microglia-centered interactions in aged tissue.

We found that the *corpus callosum*/*anterior commissure* (CC/ACO) region in the aged brain is dominated by oligodendrocyte-microglia and oligodendrocyte-oligodendrocyte crosstalk [43], while the *ventricular niche* (VEN) shifts from NSC-neuroblast-ependymal interactions in young mice to a pro-inflammatory ependymal-ependymal interaction pattern along aging, characterized by upregulation of interferon-stimulated genes (*Ifitm3, Stat1, Bst2* ; Supplementary Fig. 8c).

Focusing on oligodendrocyte-microglia interactions, we identified the genes that are upregulated in cells participating in these spatial edges (Methods 8) using the following approach: for each cell type, we compared cells that participate in high-confidence oligodendrocyte-microglia edges (“interacting”) to cells of the same type that do not (“bystander”), and performed Wilcoxon rank-sum differential expression analysis between the two groups (Methods 4.10, Supplementary Fig. 8a,b). This analysis reveals that interacting microglia upregulate *Lyz2* (Log2FC 2.308), a marker of activated myeloid cells consistent with a shift toward a phagocytic state engaged in clearing accumulating myelin fragments [44]. Our analysis further revealed that this interaction is characterized by a shift toward a pro-inflammatory, antigen-presenting phenotype, indicated by the enrichment of *B2m* (Log2FC 1.213) and *H2-K1* (Log2FC 1.392), as previously suggested [45] (Supplementary Fig. 8a). Using Casei, we also observed that interacting oligodendrocytes upregulate the complement component gene C4b (Log2FC 2.749) previously associated with age-related synaptic loss and cognitive decline [46], the stress-responsive lipocalin Apod (Log2FC 1.109), a marker of oligodendrocyte stress and neuroprotection in aging white matter [47, 48], and the serine protease Klk6 (Log2FC 1.915), associated with myelin turnover [49](Supplementary Fig. 8b).

Beyond individual interactions, we used our framework to identify coordinated gene expression programs that organize the spatial architecture of aged versus young brain tissue, through the eigendecomposition of the differential interaction matrix (Methods 4.9). The aged-specific program was primarily characterized by the cellular response to type I interferon (Fig. 5i), consistent with interferon’s established role in age-related cognitive decline and neural stem cell quiescence [50–52]. In contrast, the young-specific program was significantly enriched for the gene ontology terms *positive regulation of peptidyl-tyrosine phosphorylation, positive regulation of intracellular signal transduction*, and *gliogenesis* (Supplementary Fig. 8). These findings indicate that the spatial architecture shifts from growth-promoting signaling in the young brain to a pro-inflammatory state in the aged brain.

Many of the condition-associated interactions identified by Casei were not recovered by statistical approaches applied directly to the spatial proximity graph, including significance overlap testing and differential *z*-score analysis between conditions (Supplementary Notes 8, Supplementary Fig. 9). Neither approach recovered key interactions including aged-associated oligodendrocyte-oligodendrocyte and oligodendrocyte-microglia interactions, or young-associated ependymal crosstalk (Supplementary Fig. 9). These interactions are detected by Casei because its confidence-based filtering operates at the level of cell states rather than cell type labels. For example, Casei identifies distinct subsets of ependymal-ependymal edges as associated with both young and aged tissues, reflecting different transcriptional contexts within the same cell type pair.

## 3 Discussion

We presented Casei, a computational framework for the identification of condition-specific cell-cell interactions in spatially-informed gene expression data. By learning a bilinear interaction term between the gene expression profiles of neighboring cell pairs, Casei assigns a training dynamics-based confidence score to each edge in the spatial cellular proximity graph that reflects the degree to which it represents a condition-associated cell-cell interaction. Methodologically, Casei contributes a general framework for comparing labeled graphs across conditions with per-edge condition-association scores and interpretable-by-construction weights, applicable beyond spatial omics to any setting where the scientific question concerns which edges, rather than which nodes, differ between conditions. We validated Casei along two complementary axes: controlled semi-synthetic spatial perturbations with ground-truth perturbed edges, where it outperforms node-level, edge-level, and graph neural network baselines, and three spatial transcriptomics datasets, where it recovers biologically interpretable patterns of tissue reorganization. Applied to diverse spatially-informed transcriptomics datasets, Casei revealed biologically interpretable spatial reorganization patterns, including the shift from endothelial to macrophage-dominated networks in atherosclerosis, disruption of hepatocyte zonation in fibrosis, and region-specific oligodendrocyte-microglia interactions in aging white matter.

We note that Casei identifies spatial co-occurrence patterns and their condition-associated reorganization, which does not necessarily reflect direct cell-cell communication, as cells may co-localize due to shared microenvironmental preferences or structural constraints rather than active signaling. Additionally, preprocessing steps such as cell segmentation may introduce artifacts, as incorrectly assigned transcripts between neighboring cells can artificially inflate interaction signals. Nevertheless, Casei remains effective under these conditions, and we expect its performance to improve further as spatial transcriptomics data quality continues to advance.

Another challenge for future work is disentangling condition-associated spatial reorganization from changes in individual cell states, for example, by integrating Casei with complementary approaches that score condition-association at the level of individual cells. While in this work we focused on pairwise interactions, in future work Casei can be extended to higher-order interactions, to enable the detection of multi-cellular niches and tissue modules.

By shifting the focus of detection from *individual cells* to the *relationship between them*, Casei reveals interaction patterns that are condition-specific. Casei can be applied across a wide range of biological settings where different conditions or samples reflect changes of tissue architecture, due to either environmental factors, external perturbations or internal reorganization. Therefore, our framework can be used to characterize the changes in the spatial makeup of tissues across conditions, uncover mechanisms underlying the spatial reorganization of cells and cell types along axes of biological variation such as development, identify region-specific signaling pathways, and highlight factors driving disease onset reflected in multicellular rewiring.

## 4 Methods

### 4.1 Rationale for the Casei framework

Condition-specific changes in tissue architecture often manifest not in individual cell states but in the interactions between neighboring cells. To capture these signals, we classify edges rather than nodes in the spatial cellular proximity graph. The bilinear form directly models gene-gene co-variation between neighbors, following from the pairwise energy-based modeling described in Section 4.5. The low-rank constraint over the bilinear form keeps the model compact and interpretable, and the training dynamics confidence-based filtering retains only cell-cell edges that are robustly associated with a given condition.

### 4.2 Algorithm

#### 1. Input

- Gene expression matrices 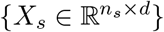, where *n*_*s*_ is the number of cells in sample *s* and *d* is the number of genes.
- Spatial coordinates 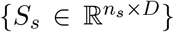, the physical location of each cell in sample *s*, where *D* ∈ {2, 3} is the number of spatial dimensions and depends on the measurement technology. Each edge (*i, j*) ∈ *E*_*s*_ is labeled with its sample’s condition *c*_*ij*_ = *y*_*s*_

#### 2. Preprocessing and Graph Construction

- Normalize gene expression: *X ←* log(10^4^ *· X*/sum(*X*) + 1).
- Construct *k*-NN spatial graphs *G*_*s*_ = (*V*_*s*_, *E*_*s*_) from coordinates *S*_*s*_.
- Label each edge with its sample’s condition *y*_*s*_.

#### 3. Bilinear Model Training

- Initialize weight matrices *W*_*c*_ ∈ ℝ^*d×r*^ for each condition *c*
- For each epoch *e* = 1 to *E*:
  – For each edge (*i, j*):
    * Compute bilinear score: 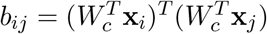.
    * Compute probability: 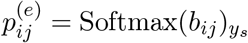
    * Store 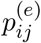 for confidence tracking.
  – Update all weight matrices 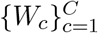 by minimizing the cross-entropy loss over all edges: 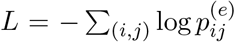 via backpropagation, with L1 (λ_L1_ = 10^*−*5^) and L2 (λ_L2_ = 10^*−*4^) regularization on *{W*_*c*_*}*.

#### 4. Confidence Computation and Filtering

- For each edge (*i, j*): compute confidence 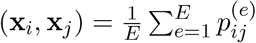 .
- Rank edges by confidence within each condition.
- Retain only the top *p*% (defaulting to 5%) of edges with the highest confidence scores for each condition. This filtered adjacency structure is referred to as the **condition-adjusted graph** (see Methods <u>4.8),</u> which is used for all downstream analyses.

#### 5. Output

- Condition-specific filtered edge sets.
- Edge confidence scores *{*confidence(**x**_*i*_, **x**_*j*_)*}*.
- Gene-gene interaction matrices 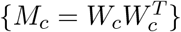.

#### 6. Downstream analysis

- Identify condition-related cellular neighborhood enrichment.
- Extract condition-related top gene pairs.
- Extract condition-associated spatial expression programs.

### 4.3 Data Preprocessing and Graph Construction

Gene expression matrices are normalized using the formula *X←* log(10^4^*· X/*sum(*X*) + 1). Spatial proximity graphs were generated by constructing *k*-nearest neighbor (*k*-NN) graphs based on cellular physical coordinates, where nodes represent cells and edges connect spatially adjacent cells. Each edge is then assigned a label *y*_*s*_ corresponding to the experimental condition of the sample from which it originated.

### 4.4 Architecture choices

We employ a bilinear architecture that explicitly captures interactions between spatially neighboring cells *i* and *j*. For gene expression profiles *x*_*i*_, *x*_*j*_ *∈* ℝ^*d*^, the interaction score is defined as:

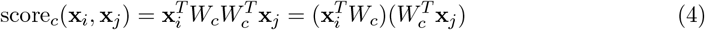

where *W*_*c*_ *∈* ℝ^*d×r*^ is a low-rank matrix (*r ≪ d*). We selected this architecture for two primary reasons:

- **Feature Interaction Modeling:** Unlike MLP-based approaches (e.g., concatenation or summation) which treat features independently prior to non-linear transformation, the bilinear layer explicitly models multiplicative gene-gene interactions essential for capturing coordinated spatial patterns [53]. While MLPs can in principle approximate such interactions, doing so requires exponentially growing model complexity compared to the quadratic scaling of explicit multiplicative architectures [54].
- **Mechanistic Interpretability:** The bilinear structure allows for direct, weight-based interpretability. The learned matrix 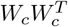 reveals specific gene pairs associated with spatial reorganization, and its symmetric nature enables eigendecomposition to extract orthogonal gene programs for functional analysis [53].

Validation on semi-synthetic datasets demonstrates that this architecture significantly outperforms MLP baselines, particularly in detecting spatial reorganization that does not alter individual cell state distributions (Fig. 2f).

To encourage sparsity in the learned interaction matrices and prevent overfitting, the crossentropy loss is augmented with L1 (*λ*_L1_ = 10^*−*5^) and L2 (*λ*_L2_ = 10^*−*4^) penalties on *W* .

### 4.5 Probabilistic modeling

We model a whole tissue as a graph, where each node *x*_*i*_ *∈* ℝ^*g*^ represents a cell with *g* genes, and nodes *i, j* are connected by an edge (*i, j*) *∈* ℰ when the spatial location of cell *j* is one of the *k*-nearest neighbors of cell *i*.

To model the effects of neighboring cells on each other, we model the energy of the whole tissue as an Ising model composed of cells interacting through the bilinear matrix:

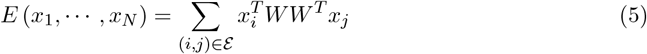

This, in turn, can be transformed to a probability assigned to the states of all cells in the tissue, given by:

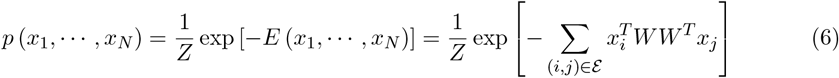

where *Z* is a normalization constant. The above is the max-entropy distribution under the constraint of pairwise interactions, and can thus be considered the simplest model for possible cross-talk between neighboring cells in a tissue.

In this work, we aim to identify which spatial cellular interactions are context-associated, namely which are enriched in specific conditions, or *cases*, for instance enriched in diseased versus healthy or in young versus old tissues. We do so by inferring interaction matrices *W* that optimally distinguish between different cases. Under this construction, and assuming for simplicity that the data contains samples of two different conditions (*c*_1_, *c*_2_) and that both conditions are a-priori equally likely, then the probability for case 1 given the cellular states in the tissue **x** = {*x*_1_, *· · ·, x*_*N*_ } is:

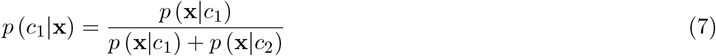

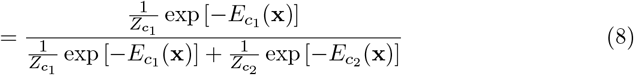

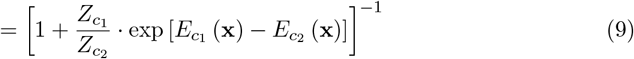

where 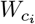 is the interaction matrix optimized for case *i*, 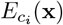 is the energy of the state **x** under the Ising model optimized for case *i*, and 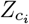 is the normalization coefficient for case *i*.

The normalizing coefficients (or partition functions)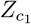 and 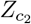 are intractable in this scenario and difficult to estimate. Instead, we will use an approximate approach. Since Equation 8 takes the form of a standard softmax function, given different samples for each case, we opt to infer 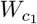 and 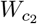 through discriminative training (Equation 2), with the standard cross-entropy loss. The interaction matrices found in this manner will in turn serve as an approximation of the true underlying probabilities *p*(**x** |*c*_1_) and *p*(**x** |*c*_2_) (through Equation 6), up to a normalization constant.

Furthermore, note that the main factor driving the classification in Equation 9 is the difference in energies, given by:

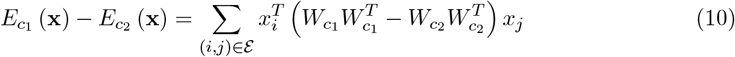

Consequently, the symmetric matrix:

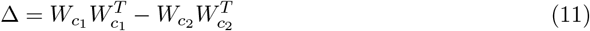

can be further analyzed to infer which specific genes underlie the differentiation between the cases. Moreover, large tissues can be broken down into smaller subsections, each modeled as a smaller Ising model, meaning that at test time, different regions within the same tissue can be individually classified.

### 4.6 Semi-Synthetic Data Generation and Benchmarking

#### Data Sampling and Graph Construction

All synthetically perturbed tissues (see 2.2) were generated based on the mouse organogenesis seqFISH dataset [17] as a biological template, accessed via Squidpy [15]. For each replicate, *n*_cells_ = 15,000 (3 sub-replicates of 5,000 cells per condition) cells were drawn with replacement from the source dataset, preserving the original expression profiles and spatial coordinates. A spatial proximity graph was then constructed using *k*-nearest neighbors (*k* = 5) applied to physical coordinates. Ten independent simulation experiments were performed per perturbation scenario.

#### Ground Truth Perturbed Edges

For all perturbation cases except for case 5, the ground truth perturbed edges are considered as *k*-NN edges incident to at least one cell that was synthetically perturbed. Case 5 defines perturbed edges directly as the union of removed and newly added edges. This classification of ground truth perturbed edges may introduce false positives, edges labeled as perturbed despite carrying no detectable signal. For example, when the modified cell’s new expression profile remains similar to its original profile or to that of its neighbor, leaving the co-expression pattern across the edge unchanged. However, it cannot introduce false negatives, since every edge carrying a detectable perturbation signal must have at least one modified cell as an endpoint, and such edges are included in our ground truth perturbed set.

This setting conservatively lowers AUROC for all estimators equally, leaving their relative ranking a valid basis for comparison.

#### Perturbation Scenarios

In Section 2.2 we presented four procedures for generating synthetic perturbations over spatial omics datasets which simulate spatial reorganization of the tissue:

- **Case 1 - Area Mixing**: Within a predefined region, cell positions are randomly permuted while their expression profiles remain unchanged. This spatial mixing disrupts the structure of the local cellular spatial graph without changing the distribution of cellular states.
- **Case 2 - Cell Type Enrichment**: A fraction *p*_enrich_ (0.5) of cells within a predefined region is replaced by cells of a predefined type. The gene expression profiles for the newly introduced cells are randomly sampled from the measured population of cells of the target type in the dataset.
- **Case 3 - Expression Profile Swapping**: For each cell of type `ct1` within a predefined region, its expression profile is swapped with that of a randomly selected neighboring cell of type `ct2`, with probability *p*_swap_ = 0.5.
- **Case 4 - Cell-Cell Interaction Enrichment**: For each cell of type `ct1` within a predefined region, each of its spatial neighbors is replaced by a randomly selected cell of type `ct2` within the same tissue region with probability *p*_neighbors_, by swapping their spatial coordinates.

All of these perturbation cases change the spatial organization of cells, while the abundances of different cell types and the distributions of their expression profiles are preserved.

Beyond the above synthetic perturbations, we considered two additional spatial reorganization cases:

- **Case 5 - Random Edge Permutation**: A fraction *p*_edges_ of existing edges are randomly selected and removed from the spatial proximity graph. An equal number of new edges are then added by randomly sampling pairs of cells that were not previously connected, directly rewiring the adjacency structure while preserving the total number of edges.
- **Case 6 - Ligand–Receptor Induction**: For each edge connecting a cell of type `ct1` to a cell of type `ct2` within a predefined region,a Poisson-sampled count *δ ∼* Poisson(*λ*_induce_) is artificially added to gene *g*_*A*_ in the `ct1` cell and gene *g*_*B*_ in the `ct2` cell. This creates a spatially localized co-expression signal present only at specific cell-cell interfaces.

#### Architectural Comparison and Benchmarking

To evaluate the predictions based on the bilinear gene-gene interactions form (see 2.2), we compared Casei against alternative architectures for combining node features into edge predictions:

- **Bilinear (C****asei**): Explicitly models pairwise interactions via 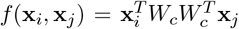, where **x**_*i*_, **x**_*j*_ *∈* ℝ^*d*^ are the gene expression profiles of the two neighboring cells, *W*_*c*_ *∈* ℝ^*d×r*^ is a low-rank projection matrix with rank *r* = 64, and *d* is the number of genes. A separate weight matrix *W*_*c*_ is learned for each condition *c ∈* {1, …, *C*}, yielding a condition-specific interaction matrix 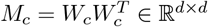.
- **Node_Mean (Baseline):** A node-level MLP classifier trained to predict the condition label from each cell’s gene expression profile **x**_*i*_ *∈* ℝ^*d*^ independently, without considering neighboring cells. Edge confidence is derived as the mean of the two incident node scores: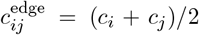. This represents the null hypothesis that edge-level perturbation detection reduces to detecting cell-level signals.
- **Graph Neural Network (GNN) baselines:** Three node-level GNN classifiers that aggregate information from spatially-proximal cells along the *k*-NN spatial proximity graph before predicting each cell’s condition. Each model consists of two graph-convolutional layers (hidden dimension 64) followed by a linear classification head, with dropout (*p* = 0.3) applied between layers. Edge confidence is derived, as in *Node_Mean*, by averaging the predicted probabilities of the two incident nodes across training epochs. These three GNN architectures form the foundation of many spatial-omics methods, including spatial domain identification (e.g., STAGATE [21], SpaGCN [55], GraphST [23]), batch integration, and cell-cell communication inference (e.g., HoloNet [22]). The three architectures differ in their neighborhood aggregation scheme:
  – **GCN** [18]: applies symmetric-normalized spectral graph convolutions, where each node’s representation is updated as a normalized weighted average of its neighbors’ features.
  – **GAT** [19]: replaces fixed normalization with learned multi-head self-attention coefficients (4 heads in the first layer, 1 in the second), allowing the model to weigh neighbors adaptively.
  – **GraphSAGE** [20]: uses an inductive mean-aggregation scheme that concatenates a node’s own features with the mean of its neighbors’ features.
- **MLP-based variants:** Three multilayer perceptron (MLP) architectures (3 fully connected layers with ReLU activations) that differ only in how the feature vectors of the two nodes are aggregated before being passed through the network: element-wise sum *f*(**x**_*i*_ + **x**_*j*_) (**MLP_Sum**), concatenation *f*([**x**_*i*_∥**x**_*j*_]) (**MLP_Concat**), and element-wise product *f*(**x**_*i*_ ⊙ **x**_*j*_) (**MLP_Product**).

All models were trained using the same hyperparameters (Adam optimizer, learning rate 10^*−*3^, batch size 512, 100 epochs). Confidence scores were computed as the mean predicted probability for the input label across all training epochs following [9]. Performance was evaluated using the Area Under the Receiver Operating Characteristic curve (AUROC) for edge-level perturbation detection, averaged over four simulation replicates for each type of synthetic perturbation.

### 4.7 Extracting gene pairs underlying condition-associated spatial reorganization

To infer the gene pairs expressed in neighboring cells associated with reorganization, we begin by computing the differential interaction matrix

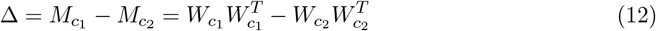

which captures changes in gene-gene relationships of neighboring cells between the test and reference conditions. When comparing against multiple reference conditions 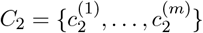, the averaged reference interaction matrix is computed as 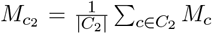. The differen-tial interaction matrix Δ contains both positive values (interactions strengthened in *c*_1_) and negative values (interactions weakened in *c*_1_).

To focus on the condition-specific programs enhanced in *c*_1_, we extract the positive component of the differential matrix as Δ^+^ = max(Δ, 0), where the operator max is taken elementwise.The top condition-specific gene pairs are identified as those with the largest entries 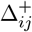 in this matrix.

To visualize these relationships, we construct a weighted gene interaction network where nodes represent genes, edges (each one represents pairs of genes) connect the top *n* gene pairs (default *n* = 10), with edge weight set to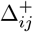, reflecting the strength of their condition-association co-expression across neighboring cells.

### 4.8 Condition-Adjusted Graph Construction

After training, each edge in the spatial proximity graph is assigned a confidence score reflecting how associated its co-expression pattern is with a given condition based on features of its training dynamics (see Methods 4.2). Edge confidence scores are stored in a confidence matrix over all cell pairs. For each condition *c*, we retain the top *p*% (default *p* = 5) of edges ranked by confidence, yielding a *condition-adjusted graph*, which is a subgraph of the original spatial proximity graph that reflects condition-associated interactions. The number of high-confidence edges incident to each cell is recorded per cell as a summary statistic, enabling downstream identification of spatially condition-associated regions. The *condition-adjusted graph* is used as input to all subsequent neighborhood enrichment analyses (see 2.3, 2.4, 2.5).

### Interaction programs

We perform spectral decomposition of the symmetric matrix Δ^+^ (the positive component of the differential interaction matrix, see Methods 4.7) to extract orthogonal gene programs. Specifically, we compute the eigendecomposition Δ^+^ = *V* Λ*V* ^*T*^, where Λ = diag(*λ*_1_, …, *λ*_*d*_) is a diagonal matrix of real eigenvalues (guaranteed by the symmetry of Δ^+^) sorted in descending order of magnitude |*λ*_1_| *≥· · ·≥*|*λ*_*d*_|, and *V* = [**v**_1_| *· · ·* | **v**_*d*_] *∈* ℝ^*d×d*^ is an orthogonal matrix whose columns **v**_*k*_*∈* ℝ^*d*^ are the corresponding eigenvectors. Each eigenvector 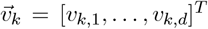 defines a gene program, where the magnitude of component *v*_*k,i*_ represents the contribution (loading) of gene *g*_*i*_ to program *k*.

#### Gene Signature Extraction

For each of the top *K* eigenvectors in *V*, we extract gene signatures by ranking genes according to their absolute loading values |*v*_*k,i*_| in descending order. The top *n* genes with highest absolute loadings define the gene signature 𝒢_*k*_ = { *g*_*i*_ : |*v*_*k,i*_| ranks in top-*n*} for program *k*. We retain both the gene identities and their corresponding loading weights {(*g*_*i*_, *v*_*k,i*_) } for downstream interpretation. To assign biological meaning to each program, we perform functional enrichment analysis against Gene Ontology Biological Processes 2023 [56] and KEGG 2021 [57] using the hypergeometric test, applying Benjamini-Hochberg FDR correction and retaining pathways with adjusted *p <* 0.05.

### 4.10 Identification of Interaction-Associated Gene Expression

To identify genes whose expression is upregulated in cells participating in a given cell-cell interaction type, we classified cells within each condition as either “Interacting” or “Bystander” based on the condition-adjusted spatial interaction network. For a given cell type pair (`ct1– ct2`), a cell of type `ct1` was labeled “Interacting” if it had at least one high-confidence edge to a cell of type `ct2` in the condition-adjusted graph, and “Bystander” otherwise.

We then performed differential expression analysis (Wilcoxon rank-sum test) comparing Interacting versus Bystander cells within each cell type, requiring a minimum of 3 cells per group.

### 4.11 Confidence Threshold Selection

The top-*p*% threshold controls the selectivity of condition-association in the resulting adjusted graph: lower values yield a sparser subgraph enriched for strongly condition-associated edges, while higher values produce a denser graph that may incorporate more ambiguous interactions. In practice, we recommend interpreting edge confidence as a relative ranking rather than an absolute score, as the magnitude of confidence values depends on factors such as dataset size and class balance. The benchmarking evaluation in Section 2.2 is performed using AUROC, a rank-based metric that is threshold-independent by construction, and therefore reflects the intrinsic quality of the confidence ranking regardless of any specific threshold value. For downstream analyses that do depend on the threshold, such as neighborhood enrichment, we verified robustness by varying *p* across a range of values. A supplementary sensitivity analysis of neighborhood enrichment results for the smoker versus non-smoker plaque comparison across different threshold values is provided in Supplementary Fig. 10, confirming that the key finding of a shift from macrophage-dominated interactions in non-smoker plaques to VSMC-dominated interactions in smoker plaques is robust to changes in threshold values.

### 4.12 Scalability Analysis

The dominant cost of training Casei is the bilinear score computation 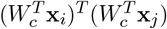 per 𝒪 edge, which requires 𝒪 (*dr*) operations through the low-rank factorization *W*_*c*_ ∈ ℝ^*d×r*^ with *r ≪ d*, avoiding the 𝒪 (*d*^2^) cost of explicitly constructing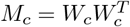. For *N* samples with on average 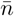 cells each and *k*-NN spatial graphs, the total number of edges scales as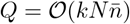.

Training over *E* epochs therefore yields an overall complexity of

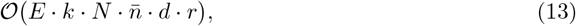

which is linear in the number of genes *d*, samples *N*, cells per sample 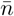, and epochs *E*. The parameter count is 𝒪 (*Cdr*) (where *C* is the number of conditions).

## 5 Data Availability

The mouse organogenesis seqFISH dataset used for semi-synthetic benchmarking is publicly available through Squidpy (https://squidpy.readthedocs.io) and the original repository [17]. The human atherosclerosis Xenium spatial transcriptomics dataset is available from [29] via GEO under accession number GSE294466. The human liver fibrosis MERFISH dataset is available from [2] via Dryad at https://doi.org/10.5061/dryad.37pvmcvsg. The mouse brain aging spatial transcriptomics atlas is available from [1] via Zenodo at https://zenodo.org/records/13883177.

## 6 Code Availability

All code for Casei is publicly available on GitHub at https://github.com/nitzanlab/casei.

## 7 Acknowledgments

We would like to thank Naomi Habib, Guy Pelc, Serafima Dubnov, and Reshef Mintz for insightful discussions.

This work was supported by The Israeli Council for Higher Education Ph.D. fellowship (J.K.), the EMBO Young Investigator Programme, the Abisch-Frenkel Foundation, and the European Union (ERC, DecodeSC, 101040660) (M.N.). Views and opinions expressed are, however, those of the author(s) only and do not necessarily reflect those of the European Union or the European Research Council.

## 8 Supplementary Notes

### Implementation Details

#### Hyperparameters

The default hyperparameters used across all experiments are summarized in Table 2.

**Table 1:** Top 10 gene pairs associated with healthy-associated cell-cell interactions in the liver fibrosis MERFISH dataset [2], extracted from the positive component of the differential interaction matrix Δ^+^ (see Methods 4.7). Gene *i* and Gene *j* denote the two genes whose coordinated expression across neighboring cells is most strongly associated with healthy tissue, and weight corresponds to the entry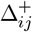.

| Rank | Gene $i$ | Gene $j$ | Weight |
| --- | --- | --- | --- |
| 1 | IL4R | LAMB1 | 0.019154 |
| 2 | A2M | KCNE1 | 0.019139 |
| 3 | CD4 | KCNE1 | 0.019066 |
| 4 | A2M | GATA4 | 0.019030 |
| 5 | A2M | IL4R | 0.019028 |
| 6 | KCNE1 | LAMB1 | 0.018946 |
| 7 | GATA4 | LAMB1 | 0.018937 |
| 8 | LAMB1 | PDGFRB | 0.018882 |
| 9 | A2M | PDGFRB | 0.018873 |
| 10 | CD4 | GATA4 | 0.018831 |

**Table 2:** Default hyperparameters of Casei.

| Hyperparameter | Value | Description |
| --- | --- | --- |
| *Graph Construction* |  |  |
| $k$ (spatial $k$-NN) | 5 | Number of spatial neighbors per cell |
| *Model Architecture* |  |  |
| Rank $r$ | 64 | Rank of bilinear factorization |
| Weight initialization | $\mathcal{N}(0, 0.001)$ | Initialization of $W$ for numerical stability |
| *Training* |  |  |
| Optimizer | Adam |  |
| Learning rate | $10^{-3}$ | |
| Batch size | 512 | Number of edges per mini-batch |
| Epochs $E$ | 100 | Total training epochs |
| Loss | Cross-entropy | Multi-class edge classification |
| *Regularization* |  |  |
| $\lambda_{L1}$ | $10^{-5}$ | L1 penalty on $W$ (sparsity) |
| $\lambda_{L2}$ | $10^{-4}$ | L2 penalty on $W$ (weight decay) |
| *Confidence Filtering* |  |  |
| Top-$p$ threshold | 5% | Fraction of highest-confidence edges retained |

#### Confidence Score Computation

At each training epoch *e*∈ { 1, …, *E*}, th e softmax probability assigned to the true label *y*_*ij*_ is recorded for every edge (*i, j*):

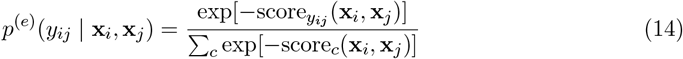

The confidence of each edge is then taken as the mean predicted probability across all epochs:

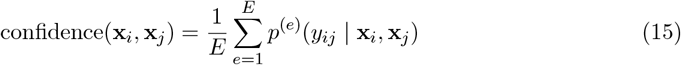

This temporal averaging reduces the influence of transient early-training fluctuations and prevents the over-confidence artifact commonly observed in neural networks trained with cross-entropy loss [9]. After training, edges are ranked by training dynamics confidence within each condition, and the top *p* = 5% are retained to form the condition-associated graph.

#### Semi-Synthetic Benchmarking Details

For the semi-synthetic analysis (Section 2.2), three replicate tissues of 15,000 cells each were generated per perturbation scenario. Gene expression for each cell was sampled from a Poisson distribution fitted to a mouse organogenesis seqFISH dataset [17]. Spatial *k*-NN graphs were constructed with *k* = 5 neighbors. Ground truth perturbed edges were defined as edges in the *k*-NN graph that have at least one incident cell that was synthetically modified by the perturbation. All baseline models (MLP_Sum, MLP_Concat, MLP_Product, Node_Mean, GCN, GAT, GraphSAGE) were trained with identical hyperparameters as Casei (Adam optimizer, learning rate 10^*−*3^, batch size 512, 100 epochs, hidden dimension 64). Performance was evaluated using the Area Under the ROC Curve (AUROC) averaged over four synthetic replicates.

#### Downstream Analysis Details

##### Neighborhood Enrichment

Condition-associated neighborhood enrichment was computed using Squidpy’s `gr.nhood_enrichment` function [15] applied to the filtered condition-specific edge sets, using 1,000 permutations.

##### Top Gene Pairs

The differential interaction matrix 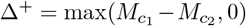 was computed based on the learned weight matrices. Gene pairs were ranked by the magnitude of their corresponding entry in Δ^+^.

##### Interaction Programs

Eigendecomposition of Δ^+^ was performed using `torch.linalg.eigh` [58]. Eigenvectors were sorted by descending eigenvalue magnitude. Gene set enrichment analysis was performed against Gene Ontology Biological Processes 2023 [56] and KEGG 2021 [57] using the hypergeometric test with Benjamini–Hochberg FDR correction (*α* = 0.05).

##### Interaction-Driven Differential Expression

Differential expression between Interacting and Bystander cells was assessed using the Wilcoxon rank-sum test, requiring a minimum group size of 3 cells. *p*-values were corrected using Benjamini–Hochberg FDR.

#### Methodological Comparison

To evaluate the utility of explicit edge modeling, we compared the performance of Casei relative to MELD [7] and Annotatability [9]. These node-centric approaches were adapted by calculating the score for each edge as the average of the scores of its two incident nodes.

As shown in Sup. Fig. 7, this averaging approach did not recover the specific hepatocyte-macrophage interaction signature in the liver fibrosis dataset (see 2.4). While node-centric methods are effective at identifying cell populations associated with a particular condition, they may lack the precision required to distinguish specific cellular crosstalk from general spatial co-occurrence.

#### Software and Hardware

All models were implemented in PyTorch [58]. Spatial graph construction was based on `sklearn.neighbors.kneighbors_graph` [58] and Squidpy [15]. Our code is publicly available at https://github.com/nitzanlab/Casei.

#### Atherosclerotic Plaques

For the analysis in Results 2.3, we used the Xenium spatial transcriptomics dataset from Pauli et al. [29], comprising 16 tissue sections from human carotid arteries: 12 advanced atherosclerotic plaque samples and 4 early lesion or control samples. Four patients contributed paired samples (both early and advanced lesions), while the remaining eight patients contributed advanced plaque samples only. Two custom Xenium gene panels were designed, each containing probes for 548 genes: Panel 1, which we used in our analysis, was designed for cell type identification based on matched scRNA-seq data, and Panel 2 captured atherosclerotic disease-associated transcripts. The dataset includes patient-level smoking history metadata for plaque samples, which we used in a secondary analysis comparing smoker and non-smoker plaques. In total, 120,164 cells were profiled across all samples after quality control and filtering (see Pauli et al. for full preprocessing details).

The plaque-adjusted neighborhood enrichment is dominated by three macrophage subtypes. MAC_HMOX1 macrophages are subluminally positioned cells primarily involved in heme degradation and lipid handling [29]. MAC_TREM2HI macrophages are lipid-associated cells located deeper in the shoulder regions of fibrous caps, implicated in extracellular matrix remodeling [29]. MAC_C1Q macrophages are characterized by their role in apoptotic cell clearance and efferocytosis [29].

The smoker-adjusted neighborhood enrichment is dominated by interactions between three VSMC (Vascular Smooth Muscle Cells) subtypes. Contractile VSMCs (VSMC_contr) represent the classical, quiescent smooth muscle phenotype maintaining arterial wall integrity [29]. Inflamed VSMCs (VSMC_inflamed) co-express classical VSMC markers alongside immune-related transcripts, representing a disease-associated phenotypic state [29]. The enrichment of interactions between and within these subtypes is consistent with previous reports of VSMC phenotypic switching in atherosclerotic lesions [59].

### Comparison with Statistical Neighborhood Enrichment Approaches

To assess whether standard spatial statistics can recover the condition-specific interactions identified by Casei in the aging mouse brain (see 2.5), we applied two complementary analyses to the original, unfiltered *k*-NN spatial proximity graphs. First, we performed Squidpy’s neighborhood enrichment independently on young and aged tissues and classified each cell type pair based on whether it reached statistical significance (BH-FDR *<* 0.05) in young only, aged only, both, or neither condition (Fig. 9c). Second, we computed a differential enrichment score by subtracting the aged *z*-score from the young *z*-score for each cell type pair, such that positive values indicate young-biased and negative values indicate aged-biased interactions (Fig. 9d).

Neither approach identified the key interactions revealed by Casei. The significance overlap analysis (Fig. 9c) classified oligodendrocyte-microglia and oligodendrocyte-oligodendrocyte as significant in both conditions, failing to distinguish it as an aging-specific interaction. The differential *z*-score analysis (Fig. 9d) was dominated by neuronal and NSC interactions, with oligodendrocyte-microglia and oligodendrocyte-oligodendrocyte pairs showing only modest differential scores that do not stand out among the many cell type pairs. The young-associated ependymal-ependymal interaction, which Casei identified, was also not identified by either statistical approach.

## 9 Supplementary Figures

**Figure 6:**
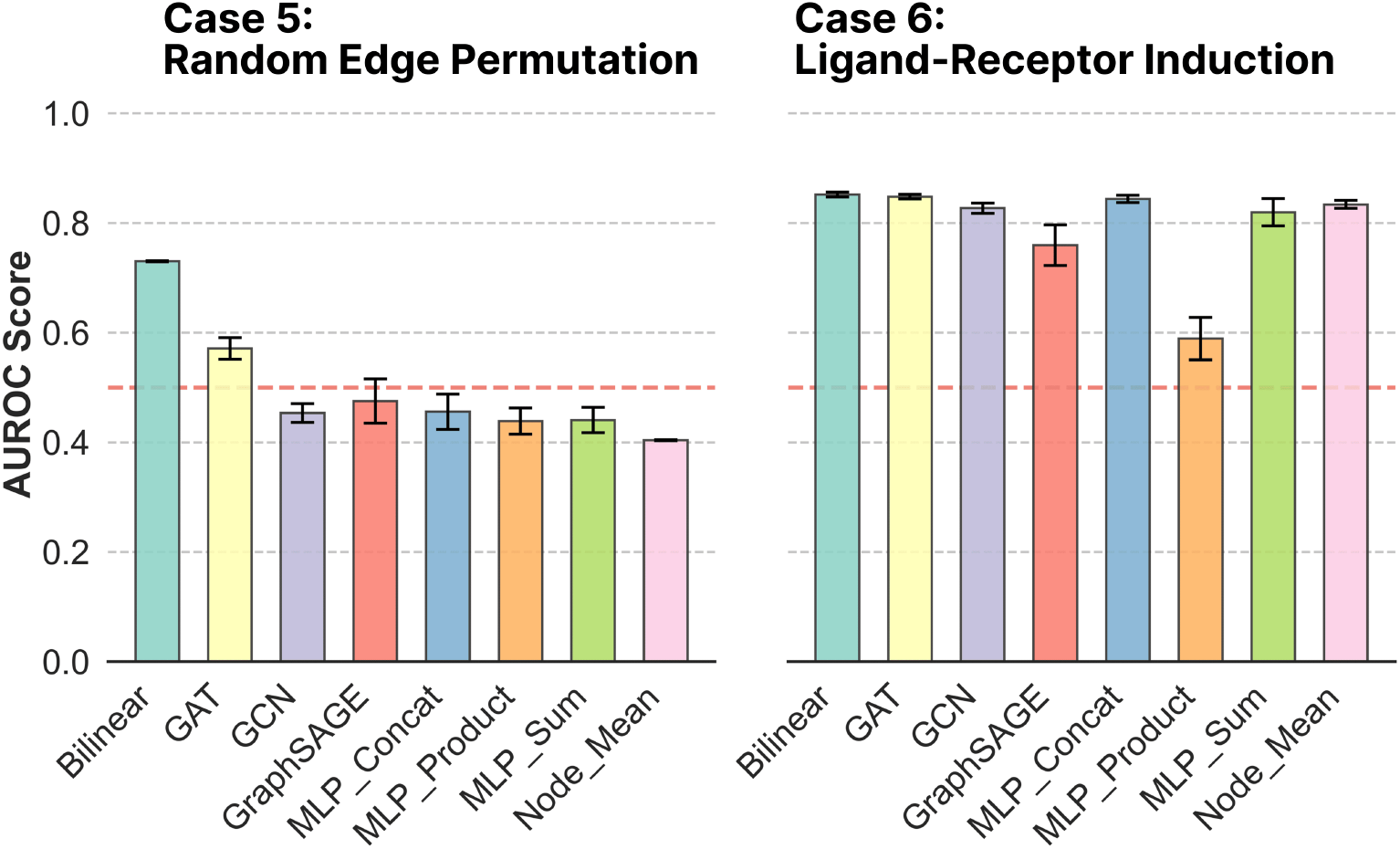
Analysis of synthetically perturbed spatial transcriptomics data. Synthetic perturbations setup based on mouse organogenesis seqFISH dataset [17]. Benchmarking performance across two perturbation scenarios. Barplots display the Area Under the ROC Curve (AUROC) for edge-level perturbation detection (*n* = 10 replicates).

**Figure 7:**
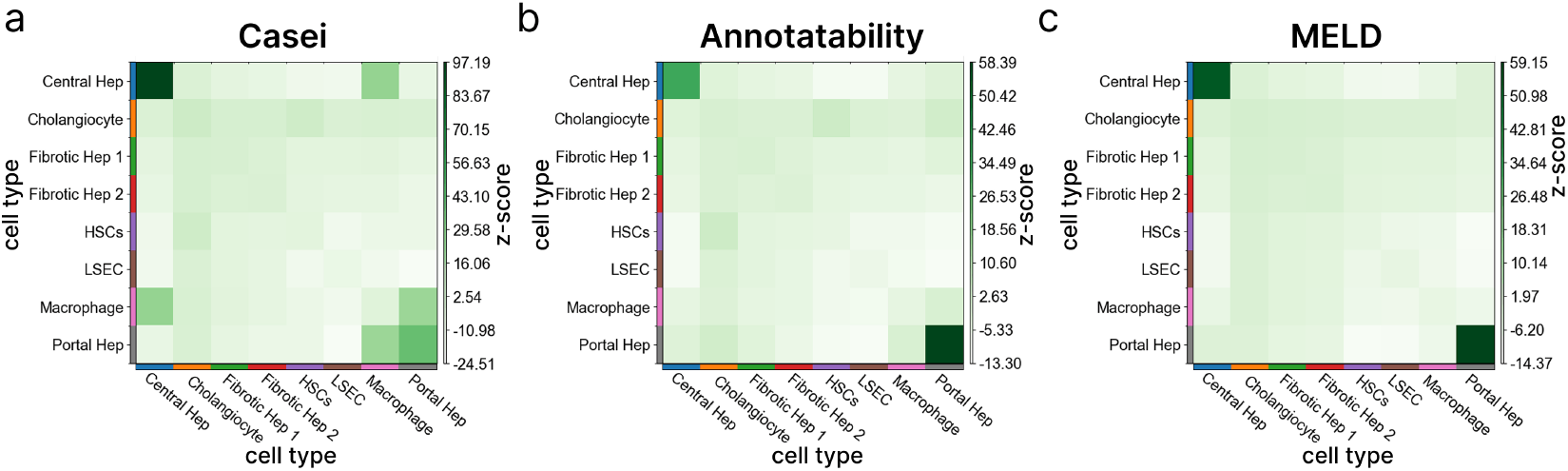
Benchmarking identification of condition-associated cell-cell interactions in liver fibrosis. (a–c) Healthy-associated neighborhood enrichment heatmaps computed on the liver fibrosis MERFISH dataset [2], comparing (a) Casei (b) Annotatability, and (c) MELD. The edge scores for Annotatability and MELD were calculated as the averaged scores of the two incident nodes (see Sup. 8).

**Figure 8:**
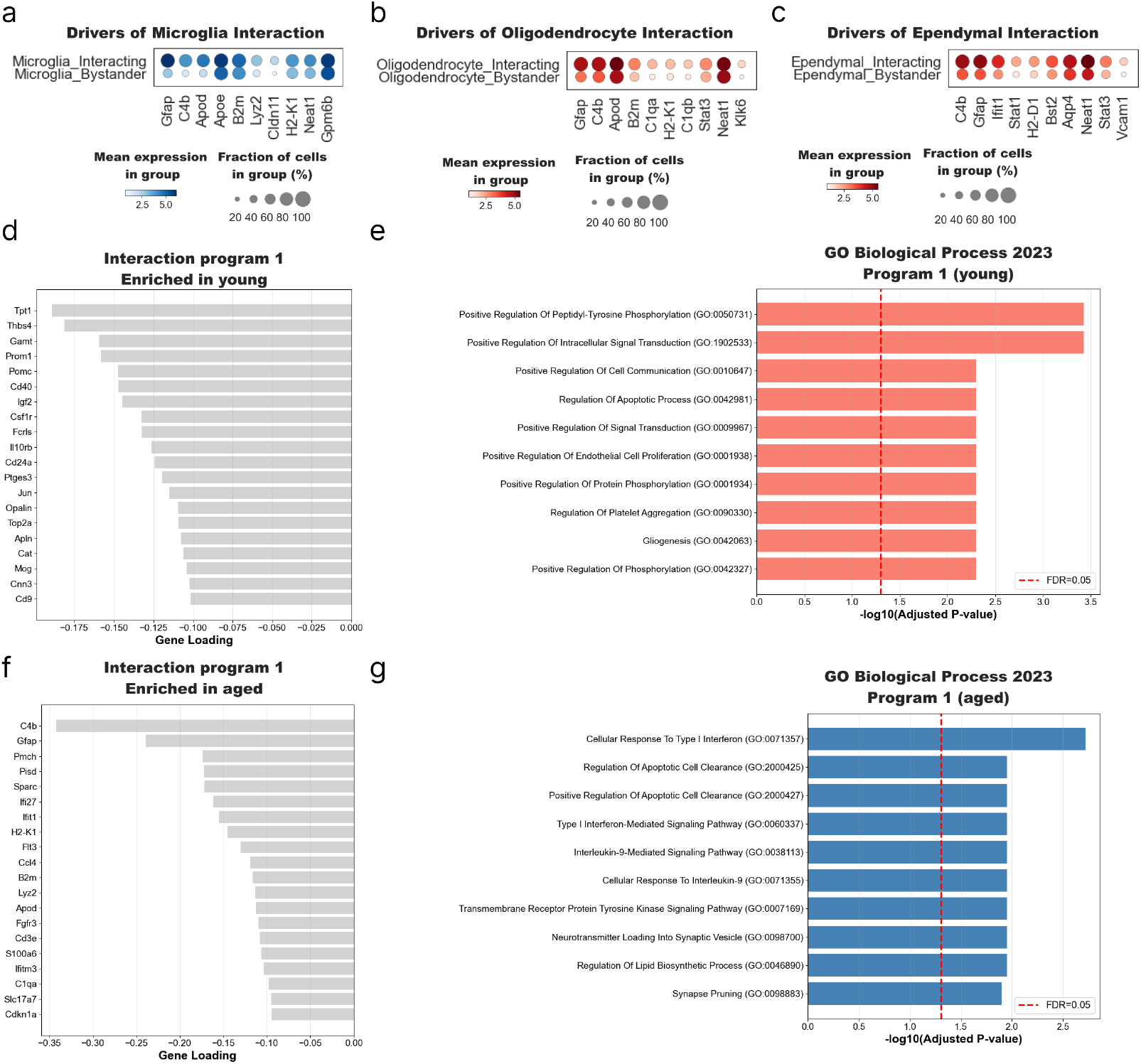
Condition-associated spatial interactions in the aging mouse brain spatial transcriptomics atlas [1]. (a–c) Dot plots comparing gene expression in cells with high-confidence spatial edges (Interacting) versus those without (Bystander) for microglia, oligodendrocytes (for microglia-oligodendrocytes interactions), and ependymal cells (for ependymal-ependymal interactions). (d-g) Characterization of the young-specific (d-e) and aged-specific (f-g) interaction programs showing top gene loadings (left) and associated GO Biological Processes (right).

**Figure 9:**
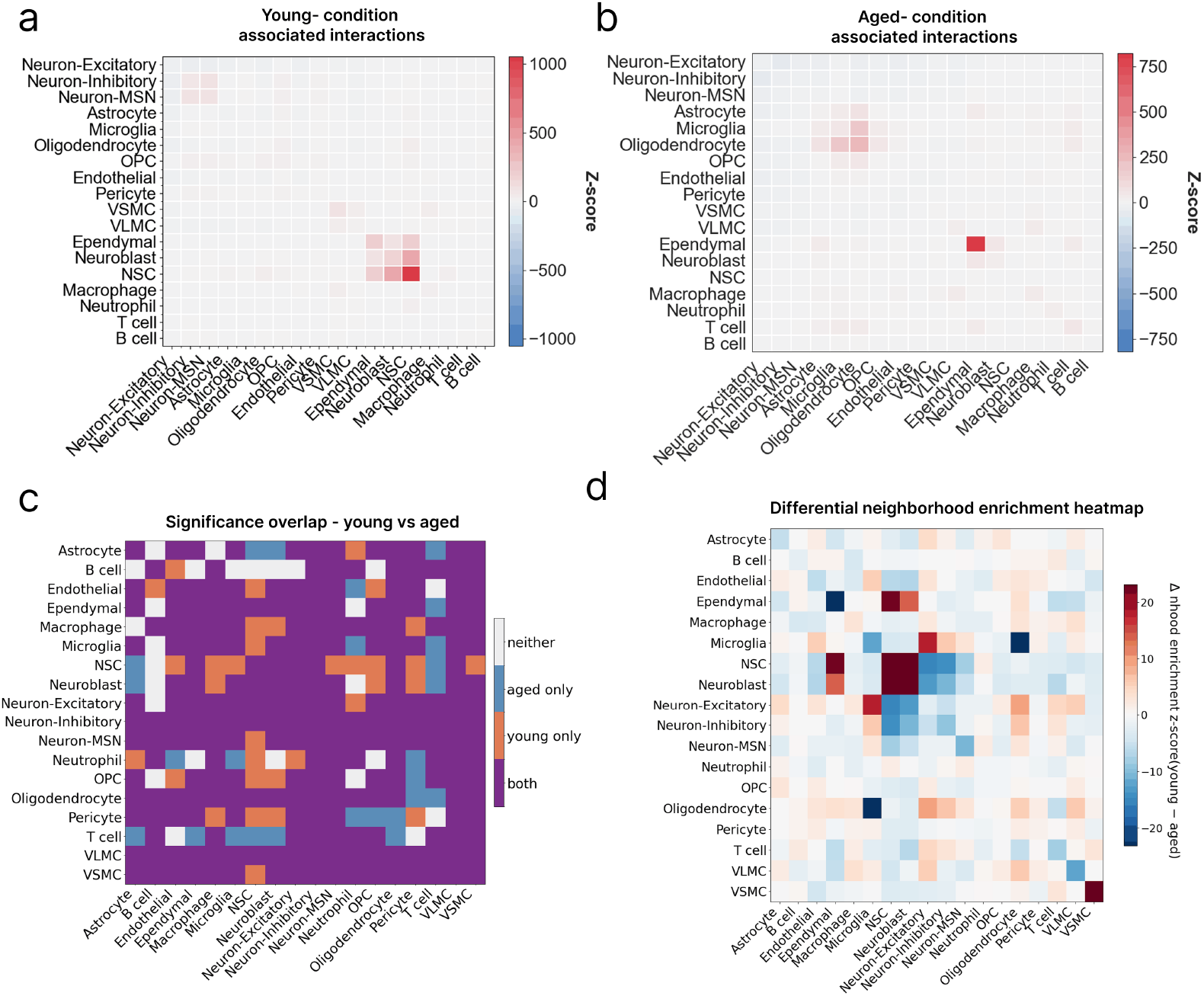
Standard spatial statistics fail to recover condition-associated cell-cell interactions identified by Casei in the aging mouse brain spatial transcriptomics atlas [1]. (a,b) Neighborhood enrichment heatmaps computed on the condition-adjusted graphs for (a) young-associated and (b) aged-associated interactions. (c) Significance overlap analysis classifying each pair of cell types by whether it is significantly enriched (BH-FDR *<* 0.05) in young only, aged only, both, or neither condition, computed over the full *k*-NN spatial proximity graph. (d) Differential neighborhood enrichment computed over the full *k*-NN spatial proximity graph. Positive (negative) values indicate young- (aged-) biased interactions.

**Figure 10:**
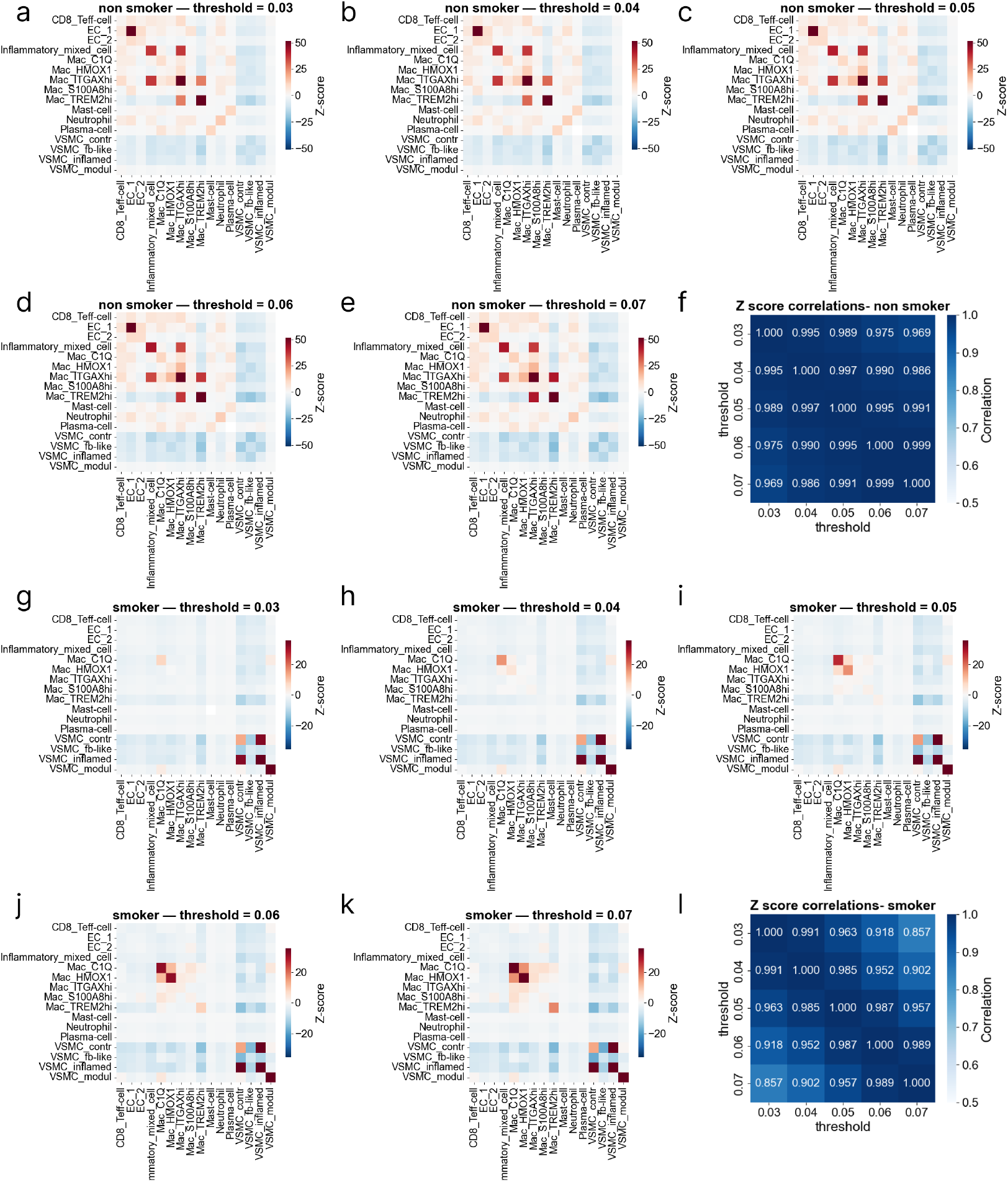
Sensitivity analysis of neighborhood enrichment relative to confidence threshold in the smoker versus non-smoker plaque comparison. Neighborhood enrichment heatmaps for non-smoker (a–e) and smoker (g–k) condition-adjusted graphs across top confidence retained edge fractions *p ∈* { 0.03, 0.04, 0.05, 0.06, 0.07} . Z-score correlation matrices across confidence thresholds for non-smoker (f) and smoker (l) conditions demonstrate robustness of the key findings: macrophage-dominated interactions in non-smoker plaques and VSMC-dominated interactions in smoker plaques.

